# Social insect colony size is correlated with rates of substitution and DNA repair gene evolution

**DOI:** 10.1101/415570

**Authors:** Benjamin E. R. Rubin

## Abstract

Larger numbers of germline cell divisions can increase the number of mutations inherited by offspring. Therefore, in systems where the number of offspring is dependent on the number of germline cell divisions, a higher overall rate of molecular evolution may be expected. Here I examine whether colony size in social insects, which varies from tens to millions, influences molecular evolutionary rates by analyzing several recently collected datasets. First, I find that colony size is negatively correlated with GC-content across 115 ant genera, indicative of a positive relationship between substitution rate and colony size. Second, genome-wide rates of molecular evolution are positively correlated with colony size in three clades of social insects including eight species in the ant genus *Pseudomyrmex*, seven fungus-growing ants, and 11 bee species. The additional germline cell divisions necessary to maintain large colony sizes might lead to mutation accumulation in the germlines of queens of these species, a process similar to that which occurs in aging human males. I also find intensified constraint on DNA repair genes in species with large colonies, suggesting that the additional mutations that occur in these taxa increase selective pressure for improved replication fidelity. Colony size, a fundamental facet of eusociality, plays a previously unappreciated role in genome evolution.

## Introduction

Rates of molecular evolution vary widely across taxa (1–3). A number of mechanisms have been identified as contributing to these rate differences, including the rate at which mutations become fixed (this can be affected by a variety of demographic effects including population size and population structure), the efficiency of DNA repair mechanisms, generation time, and the number of DNA replications in germline cells per generation. This latter trait can influence the rate of molecular evolution because gamete-producing cells accumulate mutations over an individual’s life, a process well-documented in humans; gametes produced by older males have more *de novo* mutations than younger males (4–7). Evidence for similar mechanisms has also been found in several other taxa, including birds and insects (8, 9).

The reduced effective population sizes of social insects has made them a focus for previous studies of rates of molecular evolution (9, 10). Bromham & Leys (9) compared rates of molecular evolution in pairs of eusocial and solitary taxa but found little evidence for a correlation between social behavior and evolutionary rates. While eusocial taxa, in general, do not have higher rates of substitution than solitary taxa, eusocial taxa with particularly large colony sizes (>10,000) do show the expected elevation in substitution rates (9). The authors propose that this may be due to the additional germline mutations required for queens to produce the offspring necessary to maintain these large colony sizes over their long lifespan, analogous to the process which leads to mutation accumulation in the germline of male mammals (4–7).

Here, I test the hypothesis that colony size is correlated with rates of molecular evolution. Previous studies have depended on small amounts of DNA sequence data from small numbers of taxa with little power to detect changes in rates of molecular evolution. Therefore, I examine the influence of colony size on the rate of genome-wide molecular evolution using several recently collected large-scale datasets. First, I analyze sequence data from across the ant phylogeny, testing whether colony size is related to the rates of evolution across the family. I then focus on three clades of eusocial insects that vary drastically in colony size: the fungus-growing ants, the ant genus *Pseudomyrmex*, and the bees. I leverage whole-genome sequences for taxa in each of these clades to estimate the correlation between colony size and evolutionary rate across 26 genomes. Finally, I apply genome-wide tests for negative selection to characterize differences in selective pressures on genes involved in DNA repair among species with large and small colonies. Overall, these analyses show whether any relationship exists between colony size and molecular evolutionary rate and how this connection influences the evolution of DNA repair pathways.

## Results

### GC-content is negatively correlated with colony size

Using 2,367 bp of DNA sequence data (genes *Abd-A, EF1α, LW Rh*, and *Wg*) from 115 ant genera originally collected for phylogenetic inference (11, 12), I examined the correlation between rate of molecular evolution and ant colony size (13) categorized as small (<100 individuals), medium (100-1,000 individuals), or large (>1,000 individuals). If ants with larger colony sizes have faster rates of molecular evolution then the rates of both synonymous (dS) and nonsynonymous (dN) substitutions should be higher in these taxa. However, dN would likely increase more slowly than dS due to purifying selection, leading to the expectation that dN/dS should be lower in taxa with larger colony sizes. Because there is a bias of mutations from GC to AT (14), I also investigated the association between GC-content and colony size. These traits should be negatively correlated if taxa with larger colony sizes have higher rates of mutation.

Phylogenetic generalized least squares analyses reveal that dS, dN, and dN/dS as calculated in PAML (15, 16) are not significantly correlated with colony size across 115 ant genera (dS P = 0.32, dN P = 0.076, dN/dS P = 0.36). However, I do find the expected negative correlation between GC-content and colony size, suggesting that taxa with larger colonies have higher mutation rates (P = 0.02). This appears to be driven largely by higher GC-content in the species with the smallest colony sizes (Fig. 1A; all inferred values are provided in Table S1).

**Figure 1.**
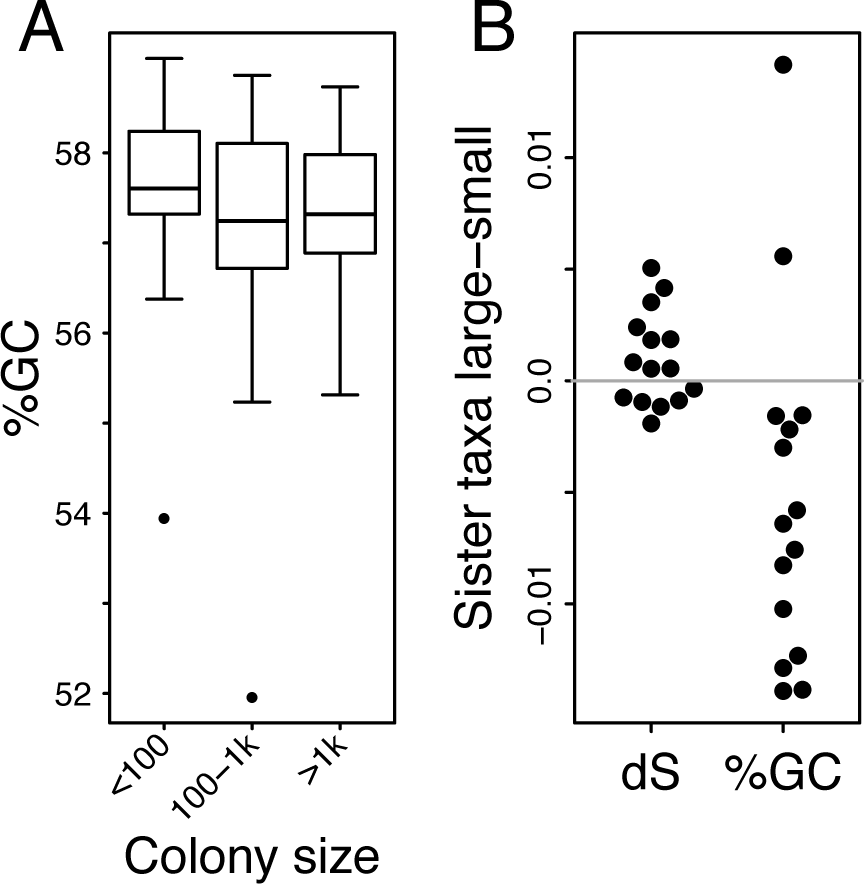
GC-content is negatively correlated with colony size across ants. A. The distribution of GC-content in four genes (*Abd-A, EF1α, LW Rh*, and *Wg*) which are conserved across ants, in three classes of colony sizes. GC-content tends to be higher in genera with smaller colonies. B. Contrasts of dS and GC-content in 15 pairs of ant sister taxa with large and small colony sizes. Both dS and GC-content show trends with colony size (positively and negatively, respectively).

To better understand these patterns and eliminate any remaining phylogenetic bias in the data, I also identified 15 pairs of sister taxa with large and small colonies (Table S2), and used paired t-tests to calculate differences in rates of evolution between these closely related taxa. These tests again yielded non-significant results for dS, dN, and dN/dS (P > 0.05) and significant differences in GC-content (t = −2.7, P = 0.02; Fig. 1B). Despite the lack of significant results for most of these metrics, trends in the expected direction are apparent. For dS, in 9/15 comparisons, large colonies have faster rates of molecular evolution than small colonies. The relatively minor differences in colony sizes between some of these sister pairs may be insufficient to generate concordant changes in evolutionary rate in this small number of highly conserved genes.

### Correlations between genome-wide rates of evolution and colony size

I hypothesized that the weak signal detected in this small number of genes across ants might be indicative of a broader genome-wide pattern, predicting that signals would be more apparent when a larger number of genomic loci were included. Therefore, I tested for correlations between rates of molecular evolution and colony size in a clade-specific fashion using whole genome data. I measured rates of molecular evolution in the orthologous groups represented by single copy genes in each taxon and tested every gene for correlations with colony size using Phylogenetically Independent Contrasts (PIC). Only a relatively small sample size (a maximum of eight in *Pseudomyrmex*, seven in the fungus-growing ants, and 11 in bees) was available for each gene. Therefore, I compiled all R^2^ values from all genes and tested whether the distribution of correlations was significantly different from zero. Although variation is expected in the strength of correlations in different genes, I predicted that the overall distribution of R^2^ across all genes would show the same patterns as above; if ants with larger colony sizes have higher rates of molecular evolution, the distributions of R^2^ for dS and dN should both be greater than zero and dN/dS should be less than zero.

I reassembled and reannotated the genomes of eight species in the genus *Pseudomyrmex* (Fig. 2A), yielding relatively complete gene sets (Table S3). These species included three that have convergently evolved large colonies (>1,000 workers) and five with small colonies (<100 workers). There were 5,260 orthologous groups of genes represented by a single copy in each taxon. As expected, the distribution of R^2^ values obtained from the PIC correlations of dS and colony size had a median of 0.054 and was significantly greater than zero (P < 1×10^−20^, Fig. 3). The dN results showed a similar signature of positive correlation with colony size (P < 1×10^−20^, median R^2^ = 0.0028, Fig. 3). In contrast but as predicted, dN/dS was negatively correlated with colony size (P = 3.01×10^−11^, median R^2^ = −0.0061, Fig. 3).

**Figure 2.**
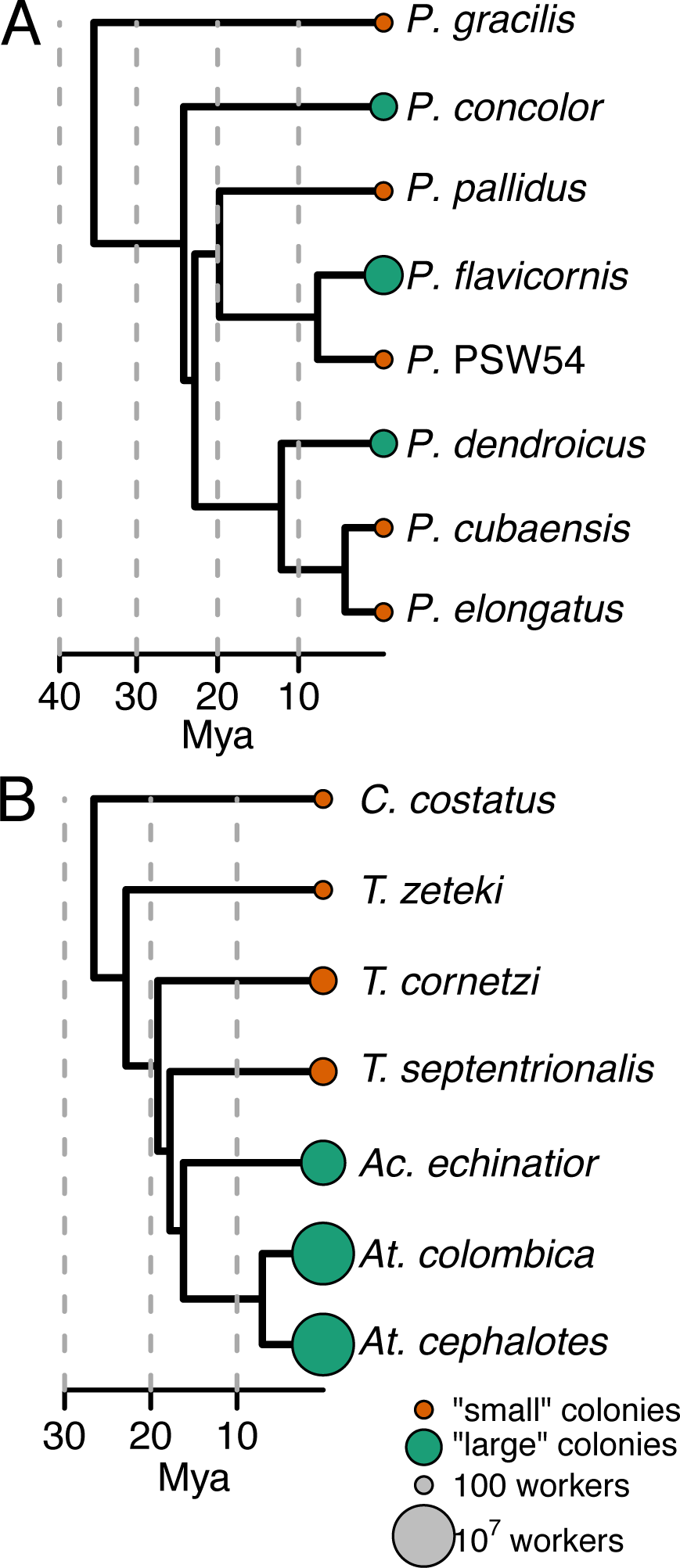
Evolutionary history of taxa used in whole-genome analyses. Phylogenies of *Pseudomyrmex* ants (A) and fungus-growing ants in the genera *Cyphomyrmex, Trachymyrmex, Acromyrmex*, and *Atta* (B). Divergence dates are from previous studies (37, 38). Colors show the taxa contrasted in tests for selection.

**Figure 3.**
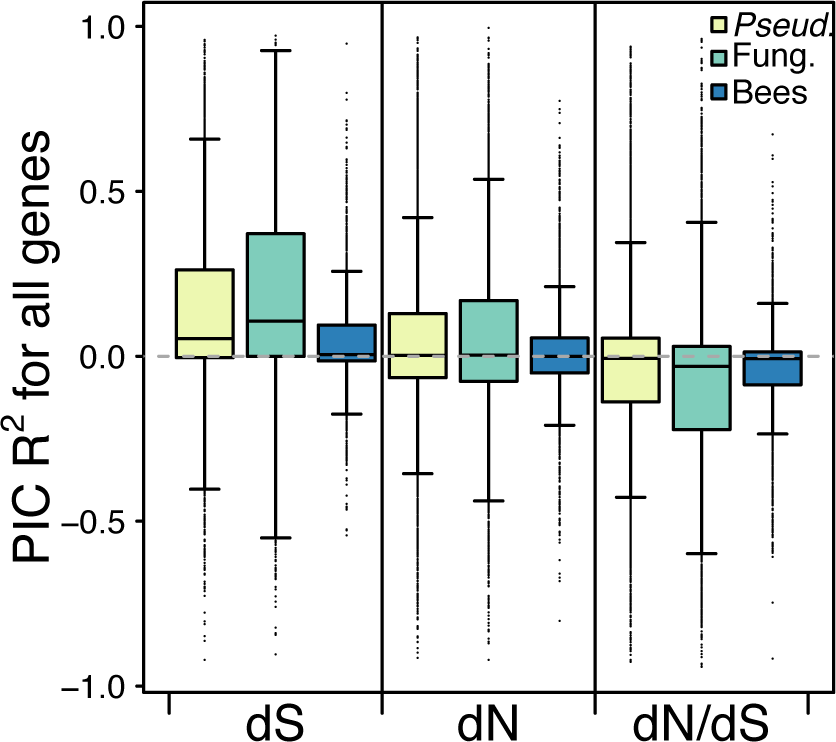
Rates of molecular evolution are positively correlated with colony size in three clades of eusocial insects. Distributions of R^2^ values of correlations between phylogenetically independent contrasts of colony size and rates of evolution for 5,260 genes in *Pseudomyrmex* (*Pseud.*), 3,980 genes in the fungus-growing ants (Fung.), and 1,975 genes in bees. Outliers are represented by dots. The distributions of R^2^ values for the correlation between dS and colony size are significantly greater than zero for all three clades as are the distributions of dN for the two ant clades. Distributions of R^2^ values for correlations between dN/dS and colony size are significantly less than zero for all three taxa.

For the seven fungus-growing ants (Fig. 2B), 3,980 genes passed quality controls and were tested for correlations between evolutionary rate and colony size. The distribution of R^2^ values for dS (median = 0.11) and dN (median = 0.0034) were both significantly greater than zero (P < 1×10^−20^, Fig. 3). For dN/dS, there was, again, a significant negative correlation between colony size and rate of molecular evolution (P < 1×10^−20^, median = −0.03, Fig. 3).

In bees, 1,975 genes were examined. The median R^2^ between dS and colony size was the lowest of any clade at 0.005 although the distribution was still significantly greater than zero (P < 1×10^−20^, Fig. 3). The distribution of correlations with dN were not significant (P = 0.05, median = 0.00005, Fig. 3). As with both ant clades, the distribution of dN/dS correlations was significantly less than zero (P < 1×10^−20^, median = −0.0068, Fig. 3). The numbers of individual genes that showed significant correlations between evolutionary rate and colony size despite small sample sizes were consistent with the expected patterns in all three clades (Supplementary Information). All of these results are robust to error in estimates of time-calibrated branch lengths (Supplementary Information).

### Signatures of selection on DNA repair

I identified genes evolving at significantly different rates in association with colony size evolution using the recently developed relative rates test (17, 18). This test examines the rate of protein sequence evolution on every branch in the phylogeny, testing for differences in the relative rates of change between the focal taxa and all other lineages.

The set of 72 genes evolving significantly slower in *Pseudomyrmex* taxa with large colonies was significantly enriched for 38 GO terms relative to all genes included in the tests (P < 0.01; Table S4). In fungus-growing ants, there were 91 GO terms significantly enriched (P < 0.01) in the 54 genes evolving significantly slower in taxa with larger colony sizes. Three GO terms were enriched in the slow-evolving genes in both of these distantly related groups of ants. These were cellular macromolecule biosynthetic process (GO:0034645), chromosome organization (GO:0051276), and DNA repair (GO:0006281). The genes assigned to each of these categories overlapped substantially, suggesting related functionality (Supplementary Information).

### Convergent constraint on DNA repair genes

One major benefit of the relative rates test is that it standardizes rates of evolution in individual genes to the overall rate of change throughout the genome, thus reducing the impact of factors that influence genome-wide rates of change. However, when characterizing purifying selection wherein rates of change are reduced, this methodology can systematically bias the results, particularly when the focal lineages are known to have higher rates of genome-wide change. If particular genes evolve at the same absolute rate in every lineage, they may be identified as evolving significantly slower in lineages where genome-wide rates of change are higher.

Therefore, I confirmed that the genes identified as evolving more slowly using the relative rates test were, indeed, evolving under additional constraint on an absolute scale. First, I estimated dN/dS ratios separately for taxa with large colony sizes and small colony sizes. For every DNA repair gene identified by the relative rates test as evolving significantly more slowly in taxa with large colony sizes, dN/dS ratios were lower in these taxa (Table S5). Second, I used time-calibrated branch lengths to calculate an absolute rate of change for every branch for each DNA repair gene and compared these rates between large and small colony taxa using Kendall’s tau correlation coefficient. For *Pseudomyrmex*, 3/8 genes were significantly more slowly evolving in species with large colony sizes (P < 0.05; Fig. S3) and for fungus-growing ants 1/7 genes was significantly more slowly evolving (P < 0.05; Fig. S4). Despite the lack of statistical significance in several of these genes, the mean rate of change was smaller in taxa with large colony sizes for every gene, indicative of an absolute increase in purifying selection.

## Discussion

In this study, I demonstrate a previously hypothesized (9) but unproven correlation between colony size and genome-wide rates of molecular evolution. Although the evidence is weak when examining small amounts of sequence data across the ant phylogeny (Fig. 1), comparing the full genome sequences of closely related species shows that taxa with larger colony sizes have higher rates of change (Fig. 3). This pattern is consistent across three distantly related clades of eusocial insects where large colonies have evolved convergently suggesting that the influence of colony size on rate of evolution is fundamental to a life history including large colonies. Remarkably, I also find increased constraint on genes involved in DNA repair in taxa with large colony sizes (Figs. S3, S4) which likely serves to compensate for the increased rate of substitution. However, given the persistent correlation between colony size and rate of molecular evolution, the consequences of increasing colony size are, apparently, not easily offset by constraint on DNA repair pathways.

This correlation is likely the result of the same mechanism responsible for increasing mutations in the gametes of aging male mammals. Mutations accumulate in germline stem cells with every additional division leading to differences in rates of substitution between species with different colony sizes. However, other processes may also contribute to the higher rate of molecular evolution in species with larger colony sizes. First, the queens of these taxa are much longer-lived than those of species with small colonies. While the accumulation of mutations in the human male germline and its impact on prezygotic mutations in offspring is well-established, the offspring of older human females accumulate more mutations during early development (19). Reproductive female ants and bees, with lifespans that sometimes stretch decades (20–22), are responsible both for the production of embryos over a long lifespan as well as continuing germline replication. In these eusocial taxa, the observed effects of aging in human males and females are likely amplified in a single sex (females). In addition, ant and bee queens only mate once, storing the collected sperm for the duration of their lives (23–26). The robustness of stored sperm in these taxa is negatively correlated with queen age (27, 28), potentially providing yet another source of mutation in long-lived queens.

A variety of taxon-specific factors might also serve to influence rates of molecular evolution in ants and bees. The number of ovarioles, each of which typically contain 2-3 germline stem cells (29), varies across species (30, 31). A larger number of ovarioles might be expected to slow overall rates of molecular evolution, as the egg-producing requirement of individual stem cells would be reduced. Although data is not available for the taxa in this study, another species of *Atta* with colony sizes in the millions (*Atta texana*) has one of the highest numbers of ovarioles known (31) suggesting that this group of ants may face selective pressures for higher ovariole number. Polygyny, a trait common in ants (25, 32, 33), could also serve a similar function as increased ovariole number, reducing reproductive load on individual queens.

Interactions between life history evolution and genome evolution have been previously identified in parasitic plants (34), flowering plants (35), and invertebrates (36) among others, although the mechanisms of these interactions are often enigmatic. However, increases in negative selection pressure on DNA repair genes concordant with increasing molecular evolutionary rates have not been reported. The implication of these two key results is that the evolution of large colony size leads to an increased rate of substitution which then leads to increased selection for DNA replication fidelity. I hypothesize that an equilibrium between these forces is eventually reached, potentially setting an upper bound on colony size and limiting the complexity of eusocial societies. Cascading effects of life history evolution on genome evolution are not often reported, particularly in eukaryotes, but are likely important factors in how genomes and gene functions change.

## Materials and Methods

### Comparisons across ants

I drew DNA sequence data and the ant phylogeny from previous studies (11, 12) focused on obtaining a genus-level phylogeny of the ants with limited sampling at the species level. Sequence data included in these studies were from four conserved coding genes: *Abd-A, EF1α, LW Rh*, and *Wg*. Several noncoding regions were also sequenced but were not used in the current study. A total of 2,367 bases of aligned coding sequence were included.

The free-ratios model of PAML v.4.9e (15, 16) was used to estimate dS, dN, and dN/dS. Because of the difficulty involved in accurately estimating rates of change over large evolutionary distances, I examined rates of change individually within the following subfamilies: Amblyoponinae, Dolichoderinae, Dorylinae, Formicinae, Myrmicinae, and Ponerinae. I only included the taxa that had sequence data for all loci (Fig. S1), limiting the estimates of dN and dS to 115 genera, concatenating sequence data from all genes. The dN and dS values estimated were standardized to time by dividing by previously reported time-calibrated branch lengths (12). PGLS analyses were run using these values using this phylogeny (12).

To identify weak trends that might be hidden in analyses of distantly related taxa, I also took a clade-specific approach to this analysis, identifying 15 pairs of closely related lineages (Fig. S2, Table S2) for which at least one genus had large colonies (>1,000 workers), and at least one had small colonies (<100 workers). For each of these pairs of genera, I again inferred dN and dS and standardized the estimates to time. I then tested for differences in evolutionary rates and GC-content between taxa with large and small colonies using paired t-tests.

### Genomic comparisons

There are two clades of ants for which previous studies have compared genomes of species spanning orders of magnitude of colony size: the fungus-growing ants and the genus *Pseudomyrmex*.

The fungus-growing ants with previously assembled genomes include *Atta cephalotes, Atta colombica, Acromyrmex echinatior, Trachymyrmex cornetzi, Trachymyrmex septentrionalis, Trachymyrmex zeteki*, and *Cyphomyrmex costatus*. *Atta* species have maximum colony sizes of 10 million whereas *C. costatus* and *T. zeteki* have colony sizes of up to 100, implying drastic change in colony size over just a short period of evolutionary time (∼25 My). All estimates of colony size for the fungus-growing ants and coding sequence annotations for the three *Trachymyrmex* species, *Atta colombica*, and *C. costatus* were taken from Nygaard et al. (37). The official gene set v1.2 for *Atta cephalotes*, the official gene set v3.8 for *Acromyrmex echinatior*, and the official gene set v2.2.3 for *Solenopsis invicta*, which I used as an outgroup, were obtained from the Hymenoptera Genome Database (hymenopteragenome.org).

*Pseudomyrmex* was originally sequenced in order to study the convergent evolution of mutualistic plant-ant behavior (38–40). Three clades of species within this genus have convergently evolved obligate mutualism with host plants in the genera *Tachigali, Triplaris*, and *Vachellia* (i.e., acacia). The genomes of the *Vachellia*-nesting *Pseudomyrmex flavicornis*, the *Triplaris-*nesting *Pseudomyrmex dendroicus*, and the *Tachigali*-nesting *Pseudomyrmex concolor*, as well as the genomes of four closely related non-mutualistic (generalist) species (*Pseudomyrmex elongatus, Pseudomyrmex pallidus, Pseudomyrmex gracilis*, and *Pseudomyrmex feralis*) were previously sequenced (39). I also included the sequence of another previously unreported species, the generalist *P. cubaensis*. Note that *P. feralis* was recently described (41) and was referred to as *P.* PSW-54 in the previous study (39). *Pseudomyrmex* is relevant to the current work because the shift to obligate plant-ant mutualism involves an increase in colony size. Estimates of colony size were not available for every species but those available for the generalists *P. gracilis* and *P. elongatus* both put maximum colony size at less than or equal to 100 (42, 43) so I used 100 as the maximum colony size for all generalists. The maximum colony size for *P. ferrugineus*, a *Vachellia*-nesting species closely related to *P. flavicornis*, is estimated to be 20,000-30,000 (44) so I conservatively used 20,000 as maximum colony size for this species. Although thorough characterization of colony size hasn’t been performed for mature colonies of *Tachigali*-nesting species, the largest *P. concolor* colony censused in a study of colonies nesting in saplings contained 1,104 workers (45). Therefore, I used 1,100 as the maximum colony size for *P. concolor* although the true maximum size of mature colonies is likely much closer to that of *P. flavicornis* given the other similarities in their biology. I was unable to find any estimates of colony size for *Triplaris-*nesting species so again used the extremely conservative 1,100 maximum colony size for *P. dendroicus*. Mutualists were previously found to have faster rates of molecular evolution (39), but that analysis was not done in a phylogenetically controlled way (e.g. (46)) so I reanalyzed these genomes here.

Significant recent work in bees has yielded genome assemblies from a dozen species (47–49) including solitary taxa and those with colony sizes in the thousands (48). I downloaded *Apis mellifera* OGS v. 3.2, *Eufriesea mexicana* OGS v. 1.1, *Bombus terrestris* OGS v. 1.3, *Bombus impatiens* OGS v. 1.2, *Melipona quadrifasciata* OGS v. 1.1, *Habropoda laboriosa* OGS v. 1.2, *Megachile rotundata* OGS v. 1.1, and *Lasioglossum albipes* OGS v. 5.42 (48, 50–52) from the Hymenoptera Genome Database and *Apis florea* NCBI annotation release 10, and *Ceratina calcarata* NCBI annotation release 100 from NCBI. The OGS for *Euglossa dilemma* was downloaded from http://sanramlab.org/scripts.html (47). Colony sizes for most taxa were taken from (48). For *C. calcarata* and *E. dilemma*, colony sizes of 10 were used.

### *Reassembly and reannotation of* Pseudomyrmex *genomes*

Seven *Pseudomyrmex* genomes were previously sequenced but only one, *P. gracilis*, was assembled *de novo* while the others (*P. concolor, P. dendroicus, P. elongatus, P. flavicornis, P. pallidus, P. feralis*) were assembled by mapping to the *P. gracilis* assembly. This methodology has the potential to bias results, reducing the presence of highly divergent parts of the genome in the assemblies. Therefore, I reassembled these genomes using the previously collected sequence data. Although not included in the original study, I also included new data from *P. cubaensis* that was collected in the same fashion as the others (half lane of 100bp paired-end sequencing on the Illumina HiSeq2000). Sequence reads were first trimmed using Trimmomatic v0.36 using the settings ILLUMINACLIP:TruSeq2-PE.fa:2:30:10 LEADING:3 TRAILING:3 SLIDINGWINDOW:4:15 MINLEN:72. I then used IDBA (53) to assemble these genomes with precorrection, a minimum kmer size of 20 and a maximum kmer size of 90. Genome completeness was assessed using BUSCO v2.0 (54) with the 4,415 conserved Hymenoptera genes in ODB9 and using the *Camponotus floridanus* AUGUSTUS profile (55).

I used RNAseq data to assemble a *de novo* transcriptome for *P. gracilis* using Trinity (56, 57) with the trimmomatic and normalize options. I also mapped these data to the genome using HISAT2 (58) with a maximum intron length of 100,000 and performed genome-guided assembly using Trinity. These two versions of the transcriptome were then combined using the PASA v2.0.2 (59) and Transdecoder pipelines (57). The mapped RNAseq data were also used to infer gene annotations directly using BRAKER v2.1.0 (60) on the repeat-masked assembly of *P. gracilis* generated previously (39).

These assembled transcripts and gene predictions were then synthesized with MAKER v3 (61, 62) in two iterations. The EST evidence passed to maker was the output from PASA, along with the gff file of genomic coordinates that PASA identified for these transcripts. The UniProt protein database, the proteins in *Apis mellifera* OGS v3.2, the proteins from *Ooceraea biroi* OGS v1.8.6, the proteins from *Cardiocondyla obscurior* OGS v1.4, and the protein sequences inferred by Transdecoder from the *P. gracilis* transcriptome were also passed to MAKER. The output from BRAKER was passed to pred_gff for this first iteration. I also used the EVM, “correct_est_fusion”, and “always_complete” options of MAKER. The previously generated custom repeat library (39) was passed for repeat masking and all other options were left as defaults. The genes without transcript or protein evidence that were predicted by this first round were checked for evidence of protein domains using InterProScan v5.21 (63) and any that showed signatures of known domains were then passed to MAKER to be included in the final annotation.

The protein sequences that were produced using this procedure for *P. gracilis* were then used to annotate the other newly assembled *Pseudomyrmex* genomes. I passed these sequences to BRAKER, using GenomeThreader to align them and train AUGUSTUS, outputting *ab initio* predictions from AUGUSTUS as well as those predictions with evidence. I then passed these results to MAKER using the same parameters as for the *P. gracilis* assembly, with the addition of the *P. gracilis* protein sequences to the protein evidence. I used the custom repeat library from the *P. gracilis* assembly for masking all genomes.

The reannotation of the *P. gracilis* genome yielded a highly complete gene set that included complete single copies of 94.9% of 4,415 conserved genes in Hymenoptera (Table S3) as identified by BUSCO (54). Using IDBA (53), I generated relatively high-quality genome assemblies from just single paired-end shotgun sequencing libraries with N50 sizes ranging from 14 kb to 36 kb (Table S3). While annotations of these genomes are clearly less complete than for *P. gracilis*, they all include more than 80% of conserved BUSCO genes, implying that they capture the majority of genes present in these genomes.

### Genome-wide rates of molecular evolution

I identified orthologous groups of genes within the clades of interest using Proteinortho v5.15 (64) setting minimum connectivity to 0.5. All resulting orthologous groups (OGs) with a representative sequence from every species were aligned using FSA (65) and alignments were filtered using Gblocks (66). I then used YN00 in PAML to estimate evolutionary rates between all fungus-growing ants and *S. invicta*, between *Pseudomyrmex gracilis* and all other *Pseudomyrmex* species, and between *Lasioglossum albipes* and all other bee species, requiring that a single sequence from every species was present in an OG to be included in the analysis. For each gene, I calculated phylogenetically independent contrasts (PICs) on dS, dN, and dN/dS using independent dated phylogenies for fungus-growing ants (37), *Pseudomyrmex* (38), and bees (67) with dates for *Apis florea* and *Euglossa dilemma* inferred from separate works (48, 68). These PICs were correlated with PICs of the log_10_ maximum colony sizes for each species using linear models in R (69) with rate as the dependent variable. R^2^ values were extracted and the sign of the correlation was recorded. The distributions of R^2^ were then tested for differences from zero using one sample Wilcoxon rank-sum tests.

Given the errors associated with time-calibrated phylogenies, I estimated the robustness of these tests in the two ant clades by generating 1,000 additional phylogenies with random branch lengths but identical topologies by randomly drawing from a normal distribution defined by the mean and standard deviation of age estimates at each node in the phylogenies. Each of these phylogenies was then used to again test for correlations between maximum colony size and dS, dN, and dN/dS in all genes.

### Tests for selection and GO enrichment

So as to include as many genes as possible in tests for selection, I reidentified and realigned orthologous genes in just the seven fungus-growing ants without the presence of *Solenopsis invicta.* For all orthologous groups in each clade, I estimated branch lengths using AAML, only including alignments at least 300 amino acids in length. I used RERconverge (downloaded from https://github.com/nclark-lab/RERconverge on March 12, 2018) with default settings to perform the relative rates test (17, 18), log-transforming branch lengths with a minimum branch length cutoff of 0.001. For *Pseudomyrmex*, I tested for significant differences in rates within the three mutualistic lineages relative to the rest of the phylogeny. For the fungus-growing ants, the three lineages with maximum colony sizes of at least 100,000 were tested against the rest of the phylogeny, including their ancestral branches in the test (i.e. (CCOS:0, TZET:0,(TCOR:0,(TSEP:0,(AECH:1,(ACEP:1, ACOL:1):1):1):0):0);). I used a significance cutoff of 0.05 and separated these significantly different genes into sets evolving significantly faster and significantly slower in the focal lineages.

I assigned GO terms to genes in *Pseudomyrmex gracilis* and *Atta cephalotes* using Trinotate (https://trinotate.github.io/) using the standard protocol. I then used these GO term assignments with GO-TermFinder (70) to test for enrichment in both the significantly faster-evolving and slower-evolving gene sets.

### Distinguishing relative and absolute purifying selection

To distinguish between relative increases and absolute increases in purifying selection in genes involved in DNA repair, I used two approaches. First, I used HyPhy RELAX (71) to estimate dN/dS separately for taxa with large and small colony sizes for the eight genes in *Pseudomyrmex* and seven genes in fungus-growing ants assigned the DNA repair GO term (GO:0006281) and evolving significantly slower in taxa with large colonies. Second, I used protein sequence evolution in a way similar to its use in the relative rates test. However, rather than standardizing rates of evolution on each lineage using the rates of evolution genome-wide, I standardized by time, using the branch lengths previously estimated (37, 38) and used Kendall’s tau correlation coefficient to compare these time-calibrated rates of change between the focal and background lineages.

## Acknowledgements

I thank Lindell Bromham, Paul Durst, Deren Eaton, Sarah Kocher, Wynn Meyer, Corrie Moreau, Luisa Pallares, Tom Stewart, and Benjamin Winger for providing analytical insights and valuable feedback on earlier versions of this manuscript. I thank the Kocher Lab at Princeton University for support. I was funded by postdoctoral fellowship grant no. 2018-67012-28085 from the USDA National Institute of Food and Agriculture.

